# Transcription factors SlMYB41, SlMYB92 and SlWRKY71 regulate gene expression in the tomato exodermis

**DOI:** 10.1101/2024.08.29.610307

**Authors:** Leonardo Jo, Sara Buti, Mariana A. S. Artur, Rianne M.C. Kluck, Alex Cantó-Pastor, Siobhán M. Brady, Kaisa Kajala

**Affiliations:** Experimental & Computational Plant Development, Institute of Environmental Biology, Utrecht University, Utrecht, the Netherlands; Laboratory of Plant Physiology, Wageningen Seed Science Centre, Wageningen University and Research, Wageningen 6708PB, the Netherlands; Department of Plant Biology and Genome Center, University of California, Davis, Davis, CA, USA; Department of Molecular, Cellular and Developmental Biology, Faculty of Arts and Sciences, Yale University, New Haven, CT, USA; Howard Hughes Medical Institute, University of California, Davis, Davis, CA USA

**Keywords:** Suberin, MYBs, WRKYs, exodermis, trans-activation assays

## Abstract

Root barrier cell types, like the endodermis and exodermis, are crucial for plant acclimation to environmental stresses. Deposition of suberin, a hydrophobic polymer, in these cell layers restricts the movement of molecules and plays a vital role in stress responses. This study investigates the role of SlMYB41, SlMYB92 and SlWRKY71 transcription factors (TFs) in regulating suberin biosynthesis in the tomato (*Solanum lycopersicum*) root exodermis by genetic perturbation. Genetic perturbation of these TFs altered exodermal suberin deposition patterns, indicating the SlMYBs as positive and SlWRKY71 negative regulators of suberization. RNA sequencing revealed a significant overlap between differentially expressed genes regulated by these TFs, suggesting a shared regulatory network. Gene set enrichment analyses highlighted their role in lipid and suberin biosynthesis as well as overrepresentation of exodermis-enriched transcripts. Furthermore, transactivation assays demonstrated that these two MYBs promote the expression of suberin-related genes, while SlWRKY71 represses them. These results indicate a complex antagonistic relationship, advancing our understanding of the regulatory mechanisms controlling exodermis suberization in tomato roots.

**Highlight:** MYB and WRKY transcription factors collaboratively regulate suberin biosynthesis in the tomato root exodermis. Antagonistic interactions may fine-tune suberization or act as a break to stop overaccumulation.

## Introduction

Root barrier cell types emerged as an evolutionary innovation that allows plants to interact and acclimate to their local environment. The endodermis and exodermis are known as two very important root barrier cell types that control the movement of molecules and solutes into the roots and their exudation into the plant’s environment (Liu and Kreszies, 2023). The endodermis is the innermost ground tissue cell layer localized next to the pericycle, and it is found in all vascular plants. It is characterized by the deposition of two hydrophobic barriers, the lignified Casparian strip and the suberin lamellae, which restrict the diffusion of molecules between the root vasculature and the surrounding environment (Robbins *et al*., 2014; Barberon *et al*., 2016; Doblas *et al*., 2017). The Casparian strip is deposited in the anticlinal walls of endodermis cells and provides a barrier for the apoplastic transport of ions and solutes, while suberin lamellae are deposited outside the plasma membrane just below the primary cell wall and prevent diffusion through the plasma membrane (Robbins *et al*., 2014; Shukla and Barberon, 2021). Because of the deposition of these hydrophobic barriers, the movement of molecules such as ions, hormones, nutrients and immunosignals are constricted to pass through the endodermis cytoplasm. Therefore, the endodermis cell layer has been proposed to act as signaling hub for root cell communication (Miyashima and Nakajima, 2011; Robbins *et al*., 2014; Salas-González *et al*., 2021; Xu *et al*., 2022b; Verbon *et al*., 2023). The exodermis is the outermost cortex layer found beneath the epidermis. Similar to the endodermis, exodermal barriers are usually composed of lignin and suberin (Barberon, 2017; Liu and Kreszies, 2023). Due to the deposition of these hydrophobic polymers, the exodermis also is an important barrier cell type in the root system, and its appearance was also an important step on plant adaptation to dry land (Ranathunge *et al*., 2011; Artur and Kajala, 2021; Liu and Kreszies, 2023). Exodermis suberization contributes to the plant drought response (Cantó-Pastor *et al*., 2024), and can act as a radial oxygen loss (ROL) barrier during flooding (Ejiri and Shiono, 2019). However, different from the endodermis, the exodermis is only found in the roots of some plant species (Artur and Kajala, 2021), particularly not in the model species *Arabidopsis thaliana*. For this reason, the endodermis has been the best-characterized barrier cell type in roots.

The suberization of barrier cell types is plastic under a wide range of abiotic and biotic cues such as osmotic, salinity, drought, nutrients, pathogens and the soil microbiome (Ranathunge *et al*., 2011; Barberon *et al*., 2016; Kreszies *et al*., 2020; Feng *et al*., 2022; Lu *et al*., 2022; Su *et al*., 2023; Kawa *et al*., 2024). Recent genetic and molecular research has identified major players of suberin biosynthesis and polymerization in response to developmental and environmental cues.

MYB (Myeloblastosis) transcription factors (TFs) have been identified as major coordinators of the expression of suberin biosynthesis genes in the Arabidopsis root endodermis (Shukla *et al*., 2021; Xu *et al*., 2022b) and seed coat (Gou *et al*., 2017). *AtMYB41* was found to induce the ectopic expression of suberin biosynthesis genes and deposition of suberin-like lamellae in multiple plant species and cell types (Kosma *et al*., 2014). *AtMYB39* and *AtMYB92* have also been shown to enhance suberin lamellae deposition when heterologously expressed in *Nicotiana benthamiana* leaves (Cohen *et al*., 2020; To *et al*., 2020). MYBs have also been shown to form multiple transcriptional sub-regulatory networks to regulate suberization of the Arabidopsis endodermis (Xu *et al*., 2022b). For example, Shukla *et al*., (2021) identified the requirement of four MYB TFs (*AtMYB41*, *AtMYB53*, *AtMYB92* and *AtMYB93*) for Arabidopsis root endodermis suberization. MYBs have also recently been shown to be involved with regulation of suberin deposition in other plant species and organs, for example in rice roots (Huang *et al*., 2024), cork oak (Capote *et al*., 2018); sugar cane (Figueiredo *et al*., 2020); kiwi fruits (Wei *et al*., 2020a,b; Han *et al*., 2022); apple fruit skins (Legay *et al*., 2016; Xu *et al*., 2022a); and in grapevine roots in response to drought (Zhang *et al*., 2020). Furthermore, chemical composition and transcriptome analyses of tomato and russet apple fruit surfaces showed a conserved gene expression signature for suberin polymer assembly, and that homologs of *AtMYB107* and *AtMYB9* regulate suberin deposition developmentally in fruit surfaces and Arabidopsis seeds (Lashbrooke *et al*., 2016).

Together with MYBs, WRKY TFs (named after their conserved N-terminal WRKYGQK motif) are emerging as potentially important regulators of suberin biosynthesis in Arabidopsis root endodermis. Arabidopsis *AtWRKY33* and *AtWRKY9* have been shown to be involved with suberization of root endodermis, conferring salt stress tolerance (Krishnamurthy *et al*., 2020). Still, little is known about the participation of WRKY TFs in MYB-centered gene regulatory modules regulating suberin biosynthesis in the Arabidopsis endodermis and other plant species and cell types.

Although there is increasing information about the regulation of suberin biosynthesis in different plant organs as well as the control of endodermis suberization and its physiological relevance, little is known about the molecular mechanisms controlling exodermis suberization, despite its importance for plant physiology and root plasticity. This is mainly due to the absence of an exodermis in Arabidopsis roots (Liu and Kreszies, 2023). Shiono *et al*. (2014) investigated gene expression during the formation of the rice exodermal radial oxygen loss (ROL) barrier and identified several TFs, including MYBs and WRKYs, as potential regulators of the suberin biosynthesis genes in rice exodermis. More recently, Kajala *et al*. (2021) identified exodermis-enriched gene expression patterns in tomato roots (*Solanum lycopersicum*). For example, a WRKY TF (encoded by *Solyc02g071130.3.1*) and a MYB TF (SlMYB41 encoded by *Solyc02g079280.3.1*) were identified as exodermis-enriched (Kajala *et al*., 2021). Reynoso *et al*. (2022) found that the plastic response of rice exodermis suberization and nutrient homeostasis in response to water deficit and rewatering might involve the interplay between MYBs and WRKYs in rice. Specifically, they identified a water-deficit-promoted suberin regulatory network gene promoters enriched in MYB and NAC TF binding sites (TFBSs). Conversely, they found that gene promoters related to iron homeostasis were enriched in basic helix-loop-helix (bHLH) and WRKY TFBSs and strongly downregulated during water deficit. Given that iron deficiency delays suberization in the endodermis (Barberon *et al*., 2016), the MYBs and WRKYs may modulate the opposing dynamics of suberization and nutrient uptake in the rice exodermis.

The expression of tomato *WRKY Solyc02g071130.3.1* (here referred to as *SlWRKY71* according to homology to Arabidopsis *AtWRKY71* (Karkute *et al*., 2018; Wang *et al*., 2020b; Kumar *et al*., 2023) has also been shown to change in response to drought stress in leaves (Karkute *et al*., 2018), and in response to jasmonate and aluminum treatments in roots (Wang *et al*., 2020b), which suggest a role in stress-responsive tomato environmental plasticity. Furthermore, recent research has shown that SlWRKY71 works as an inhibitor of gene expression to negatively regulate tomato fruit ripening (Sun *et al*., 2023).

Recently, a set of MYB TFs (SlMYB41, SlMYB92, encoded by *Solyc05g051550.2.1,* and SlMYB63 encoded by *Solyc10g005550.3.1*) were shown to be necessary for suberin deposition of the tomato root exodermis, and the suberization of the exodermis contributing to the tomato drought response (Cantó-Pastor *et al*., 2024). Whether MYBs and WRKYs act together to form a regulatory module to control the expression of suberin biosynthesis genes in tomato root exodermis is yet to be explored.

To address this question, we investigated if SlMYB41, SlMYB92 and SlWRKY71 control the expression of suberin biosynthesis genes using overexpression (OX) and CRISPR-Cas9 knock- out (ko) mutants in tomato hairy root lines (Ron *et al*., 2014). As previously reported (Cantó-Pastor *et al*., 2024), we observed a reduction in suberin in *Slmyb41-*ko and *Slmyb92-*ko lines, while their overexpression did not alter the observed patterns. In contrast, we found that the exodermis suberin deposition was reduced in *SlWRKY71*-OX. We performed RNA-seq analysis on root tips from overexpression and knockout lines to identify genes regulated by SlMYB92, SlMYB41, and SlWRKY71. Knockout of these TFs significantly impacted the root tip transcriptome, with SlMYB92 and SlMYB41 showing overlapping gene regulation, particularly in lipid biosynthesis and oxidative stress pathways. Our transcriptome findings also suggested that SlMYB92 and SlMYB41 act as positive regulators, while SlWRKY71 serves as a repressor of suberin biosynthesis genes in the tomato root exodermis. Transactivation assays further confirmed that SlWRKY71 antagonizes the promoter activation by SlMYBs. Together, our findings highlight an interplay between these three TFs to fine-tune the expression of suberin biosynthesis gene in the tomato root exodermis.

## Methods

### Phylogenetic tree construction

We retrieved the AGI locus codes and nomenclature of Arabidopsis MYBs with described functions associated with suberization (Kosma *et al*., 2014; Du *et al*., 2015; Lashbrooke *et al*., 2016; Gou *et al*., 2017; Cohen *et al*., 2020; Shukla *et al*., 2021; Xu *et al*., 2022b) from Dubos *et al*. (2010). For WRKYs, we retrieved AGI codes and nomenclature of Arabidopsis WRKY family members described by Eulgem *et al*. (2000) and Wu (2005) and we searched in the literature for the ones with described functions associated with suberization in multiple plant species (Supplementary Table S1). Using the gene identifiers we queried the primary protein sequences of MYBs and WRKYs in Phytozome 13 (https://phytozome-next.jgi.doe.gov/) using sequences from *Arabidopsis thaliana* genome TAIR10 (Goodstein *et al*., 2012). For each TF family, we performed multiple sequence alignment using MAFFT online server (https://mafft.cbrc.jp/alignment/server/) using the standard automated method with the L-INS-i model. We used IQ-TREE webserver (http://iqtree.cibiv.univie.ac.at/) to infer a Maximum Likelihood tree with 1,000 bootstraps, and we edited the phylogenetic tree using iTOL (Letunic and Bork, 2016).

### DNA constructs for gene editing and overexpression

We performed transcriptional reporter construct cloning and imaging as described in Kajala *et al*. (2021) and tomato CRISPR-Cas9 construct cloning, mutant line generation, and analyses according to Cantó-Pastor *et al*. (2024). We generated the overexpression lines by retrieving the coding sequence (CDS) of *SlMYB41* (*Solyc02g079280*), *SlMYB92* (*Solyc05g051550.2.1*) and *SlWRKY71* (*Solyc02g071130.3.1*) from the Sol Genomics database (https://solgenomics.net; ITAG3.2). We extracted RNA and synthesized cDNA from tomato (cultivar M82) as described by Kajala *et al*. (2021). We amplified the CDS without the stop codon using primers listed in Supplementary Table S2, purified the PCR products from the agarose gel using QIAquick Gel Extraction kit and introduced the cleaned fragments into the pENTR/D-TOPO vector (Invitrogen) following manufacturer’s instructions. The previously described *35S* promoter (Ron *et al*., 2014) was cloned into the pENTR5’/TOPO (Invitrogen). We synthesized 3xFLAG oligodimer using the primers listed in Supplementary Table S2 and introduced it into the pENTR/D-TOPO vectors containing the CDSs to generate CDS-3xFLAG constructs using restriction site-based cloning. To generate the final *35S*::CDS-3xFLAG binary vectors, we recombined the promoters and CDS- 3xFLAG fusions into the Multisite Gateway vector pK7m24GW (https://gateway.psb.ugent.be/search) using LR Clonase II Enzyme mix (Invitrogen). All constructs generated were confirmed by Sanger sequencing.

For the knock-down constructs of *SlWRKY71*, target guide-RNAs were generated using the CRISPR-P (http://crispr.hzau.edu.cn/CRISPR2/) and CHOPCHOP (https://chopchop.cbu.uib.no/) web-tools. Their specificity was accessed by blasting the core sequence of the guide-RNA into the tomato ITAG4.0 genome version in phytozome (https://phytozome-next.jgi.doe.gov/). Oligos for two guide-RNAs per target (Supplementary Table S3) was synthesized and cloned into pMR217 and 218 vectors, and assembled with Gateway into binary vector pMR290 as described earlier by Cantó-Pastor *et al*. (2024) (based on Fauser *et al*., 2014; Bari *et al*., 2019). The final CRISPR constructs were then introduced to *R. rhizogenes* (strain ATCC15834) and used for the subsequent hairy-root transformation steps.

### Hairy root cultures and tomato transformation

Hairy root cultures were generated following the protocols in Ron *et al*. (2014) as follows. To generate competent *Rhizobium rhizogenes* (strain ATCC15834) cells, they were grown in MG/L media (Ron *et al*., 2014) at 28-29°C for 24-30 hours to a final optical density (OD600) of 0.5-0.7. We cooled the cells on ice for 15 minutes, spun at 500 rpm for 10 min and then resuspended them in 0.01x of the culture volume of 10% glycerol. The aliquots were then immediately frozen in liquid nitrogen and stored at -80°C until use. For transformation, we incubated *R. rhizogenes* aliquots on ice with 1 μL of the desired plasmid DNA construct prior to electroporation. Then, we immediately transferred cells to 1 mL of MG/L media and shaken at 28-29°C for 3 hours. We then plated the whole aliquot on MG/L media with agar containing the appropriate antibiotics and incubated for 3 days at 28-29°C.

To generate tomato hairy root cultures, we took the following steps: tomato seeds (*Solanum lycopersicum* cv. M82, LA3475) were sterilized with 70% ethanol for 10 minutes, rinsed with sterile MQ water, soaked in 50% commercial bleach for 10 minutes and rinsed three times with sterile MQ water. We plated the seeds on MS media supplemented with 1% sucrose and incubated under 16 h light : 8 h dark for 7-10 days. We cut the expanded cotyledons in sterile conditions with a scalpel and immediately immersed in the desired *R. rhizogenes* suspension for 20 minutes, blotted on sterile Whatman filter paper and transferred to MS plates supplemented with 3% sucrose without antibiotics for 3 days at 22-25°C in dark. After three days, we transferred the explants bottom-side up to MS media supplemented with 3% sucrose, 200 mg/L cefotaxime and the desired antibiotics for transformant selection and incubated at 22-25°C until roots started to emerge. We transferred roots of about 1 cm length on the same media, and we maintained the cultures by transferring 3-4 cm root segments onto fresh plates every 2-3 weeks. After two rounds of culturing hairy roots, we placed them onto media without antibiotics.

We validated the hairy root cultures for successful gene editing events and protein overexpression. To identify the deletions in the target genes, we amplified the genomic fragment of *SlWRKY71,* and we identified the deleted region through Sanger sequencing (Supplementary Figure S1). We validated the correct protein expression of the hairy root lines overexpressing SlMYB41, SlMYB92 and SlWRKY71 through a Western blot experiment. After extracting total protein lysates from the hairy root samples, we boiled the lysates for 1 min at 96°C and we loaded them on a 4% to 15% mini-PROTEAN TGX stain-free protein gels (Bio-Rad). We transferred the proteins onto a polyvinylidene difluoride (PVD) membrane using a Bio-Rad transblot turbo and we blocked the blots with 5% (w/v) milk powder (Elk) in Tris-buffered saline (TBS) buffer. We detected the proteins using the monoclonal antibody ANTI-FLAG M2 (Sigma no. F1804, 1:1000 in 0.5% [w/v] Elk TBS). We used the goat anti-mouse IgG conjugated with horseradish peroxidase as secondary antibody (Cell Signaling Technology no. 7076, 1:2500 in 0.5% [w/v] Elk TBS plus Tween 20 0.1% [v/v]). The labeled proteins were visualized using a 50/50 mix of chemiluminescence substrates (cat no. 34096, Thermo Fisher Scientific) on a ChemiDoc Imaging system (Bio-Rad).

### Expression maps

We created the spatial expression profiles of *SlMYB41*, *SlMYB92* and *SlWRKY71* using the R package ggPlantmap (Jo and Kajala, 2024). The expression data used for plots was obtained from the tomato translatome dataset from Kajala *et al*. (2021). Data plotted is from TRAP marker lines driven by following promoters: epidermis *AtWER*, exodermis *SlPEP*, inner cortex *AtPEP*, endodermis *SlSCR*, xylem *AtS18*, phloem *AtS32*.

### Suberin & lignin histochemistry and imaging

We prepared root cross sections of hairy roots subcultures of the control (empty vector), *35S::MYB41-3xFLAG* (*SlMYB41-*OX), *35S::MYB92-3xFLAG* (*SlMYB92-*OX), *35S::WRKY71-3xFLAG* (*SlWRKY71-*OX), *Slmyb41*-ko, *Slmyb92*-ko, *Slwrky71*-ko(4), and *Slwrky71-*ko(5) that had grown for 3 weeks at 20°C. We selected newly emerged roots from the previous subculture. Of those selected roots, we harvested 2-cm sections located in regions with newly emerging lateral roots. After harvest, we fixed the roots by vacuum infiltrating with 4% paraformaldehyde (Thermo Scientific) for 1 h. After that, we placed the roots in ClearSee (Ursache *et al*., 2018) for a minimum of three days. We embedded the cleared and harvested pieces in 3.5% agarose (Fisher Scientific) and prepared 250 µm thick sections using the VT 1000 S vibratome (Leica). To visualize suberin, we incubated the cross sections on a shaker in dark for 20 min in Fluorol Yellow 088 (Chem Cruz, Santa Cruz Biotechnology Inc.) (FY; 0.01% w/v in 96% ethanol, first dissolved in 1 ml DMSO (0.5% (w/v)). After that, we counterstained the sections using aniline blue (Sigma Aldrich) (0.5% w/v, dissolved in water) and placed them on a shaker in dark for 20 min at room temperature. We rinsed the sections with MQ water until the excess aniline blue was completely washed off and kept them in 50% glycerol until imaging. To visualize lignin, we stained the cross-sections with Basic Fuchsin (BF; 0,2% w/v, dissolved in ClearSee (Ursache *et al*., 2018)) and Calcofluor White (CW; 0,1 % w/v, dissolved in ClearSee). We first stained the sections with BF on a shaker in the dark for 20 min at room temperature. The BF was rinsed off with four washes of ClearSee of 15 min each. Afterwards, we stained the sections with CW and the excess was washed off with ClearSee. After this final wash of 30 min, we kept the stained sections in 50% glycerol until imaging. We imaged all stained samples on the same day using the Zeiss Airyscan LSM 880. Samples stained with FY were imaged using 488 nm excitation and 500-550 nm detection range. Samples stained with BF & CW were imaged using an excitation range of 561 nm and detection range of 600-650 nm for BF, and using excitation range of 405 nm and a detection range of 425– 475 nm for CW.

### RNAseq tissue harvest, library preparation and sequencing

To collect the material for RNAseq, we subcultured the hairy roots of lines of the control (empty vector), *35S::MYB41-3xFLAG* (*SlMYB41-*OX), *35S::MYB92-3xFLAG* (*SlMYB92-*OX), *35S::WRKY71-3xFLAG* (*SlWRKY71-*OX), *Slmyb41*-ko, *Slmyb92*-ko, *Slwrky71*-ko(4), and *Slwrky71-*ko(5) on sterile square Petri dishes (120x120x17 mm, Greiner Bio One) on MS medium (Duchefa Biochemie) supplemented with 3% sucrose, 200 mg/L cefotaxime sodium (Duchefa Biochemie), 1% BD Difco agar (Fisher Scientific), 0.5 g/L MES hydrate (Thermo Fisher Scientific) and grew them at 25°C in the dark. After 9 days of growth, we collected the first cm of 10 root tips for each of the three biological replicates per line and snap-froze them in liquid nitrogen. The samples were collected in 2-ml tubes containing ceramic beads and ground using a tissuelyser (Retsch MM300). The mRNA extraction and library preparation were performed according to the non-strand specific random primer-primed RNAseq library protocol of Townsley *et al*. (2015). The libraries were pair-end sequenced (50 bp) at Utrecht Sequencing Facility (Utrecht, The Netherlands) using Illumina NextSeq2000.

### RNA-seq data processing

We quality trimmed paired-end reads using Trim Galore (version 0.6.6) (Krueger, 2015) using default settings. After trimming, we pseudo-aligned the paired reads to the tomato reference transcriptome version ITAG4.1 (Sol Genomics, https://solgenomics.net) using Kallisto (version 0.46.2) (Bray *et al*., 2016) with the following parameters: -b 100. Pseudo-counts were then used for differential expression analysis. All scripts used for the trimming and alignment can be found in https://github.com/leonardojo/tomato-TF-exodermis-2024.

### Differential expression analysis

We identified differentially expressed genes (DEGs) using the R package limma (Ritchie *et al*., 2015) according to Kajala *et al*. (2021, https://github.com/leonardojo/tomato-TF-exodermis-2024). We set the threshold of adjusted P value < 0.05 for DEG identification. To obtain the DEGs, we compared each mutant and overexpressor to the empty vector control hairy roots (Supplementary Dataset S1). To identify candidate genes for transactivation assays, we compiled an overview file (Supplementary Dataset S2) where we cross-referenced our DEGs with exodermis-enriched transcripts (Kajala *et al*., 2021) and a suberin module from tomato single cell data (Cantó-Pastor *et al*., 2024). We selected genes that were exodermis-enriched, good candidates for suberin barrier formation, or DEGs in our data.

### Gene ontology (GO) enrichment analysis

We tested our DEG lists for GO term enrichment using the R package goseq (Young *et al*., 2010). GO functional annotation was extracted for ITAG4.0 from Phytozome 13 (https://phytozome-next.jgi.doe.gov/). A q-value threshold of 0.05 was used to determine the enriched GO categories in each DEG list.

### Identification of tomato suberin biosynthesis genes

To define the list of fatty-acid and suberin biosynthesis genes in tomato, we used the Tomato Plant Metabolic Network database, TOMATOCYC 4.0 (https://plantcyc.org/databases/tomatocyc/4.0), and identified the tomato genes associated with the following pathways and reactions: very long chain fatty acid biosynthesis I, very long chain fatty acid biosynthesis II, suberin monomer biosynthesis and esterified suberin biosynthesis.

### DNA constructs for trans-activation assay: Promoters

We extracted tomato genomic DNA from leaves of *Solanum lycopersicum* cultivar M82 and used it as template. We used PCRBIO HiFi Polymerase (PCR Biosystems Ltd.) to amplify the promoter regions of the genes listed in Supplementary Table S4 using the primers listed in Supplementary Table S5. The PCR products were gel-purified using the kit NucleoSpin Gel and PCR Clean-up (MACHEREY-NAGEL). We cloned all the promoters, except for *Solyc02g071130.3.1*, directly into the destination vector pDLUC15 (Jo *et al*., 2020) using the NEBuilder® HiFi DNA Assembly Cloning Kit (NEB, England) after digesting it with the enzyme FastDigest *Hind*III (Thermo Fisher Scientific). Instead, we cloned the promoter sequence of *Solyc02g071130.3.1* first into the Gateway vector pDONR207 (Thermo Fisher Scientific) using the Gateway BP clonase II enzyme mix (Thermo Fisher Scientific) and subsequently into the Gateway vector pGWL7 (VIB, Belgium) using the Gateway LR clonase II enzyme mix (Thermo Fisher Scientific). We used all the cloning reactions to transform the NEB Stable Competent *E. coli* (High Efficiency) (NEB, England). We left the transformed cells to recover for one hour at 30°C in shaking conditions and then plated them on LB plates containing the appropriate selecting antibiotic. We extracted the plasmid DNA samples from the colonies using the QIAprep® Spin Miniprep kit (Qiagen) and sequence-verified them using the Sanger method (Macrogen, The Netherlands). The sequences of the promoters cloned in this study are listed in the Supplementary Table S6.

### DNA constructs for trans-activation assay: Transcription factors

We extracted RNA from leaves of *Solanum lycopersicum* cultivar M82 using RNeasy Mini kit (Qiagen) followed by DNase I (Thermo Fisher Scientific) treatment. We synthesized cDNA using RevertAid RT Reverse Transcription Kit (Thermo Fisher Scientific) together with random hexamer primers (Thermo Fisher Scientific). We amplified the coding sequence of *SlWRKY71* using this cDNA as template based on the gene model information of version ITAG2.4 (Supplementary Figure S1), while the coding sequences of *SlMYB92* and *SlMYB41* were amplified from the plasmids pENTR/D-TOPO previously cloned for overexpression. We amplified the CDS with Phusion High-Fidelity DNA Polymerase (Thermo Fisher Scientific) and the primers listed in Supplementary Table S7. We recombined the PCR products into the Gateway vector pDONR221 (Thermo Fisher Scientific) using the Gateway BP clonase II enzyme mix (Thermo Fisher Scientific) and subsequently into the Gateway vector p2GW7 (Karimi *et al*., 2002) using the Gateway LR clonase II enzyme mix (Thermo Fisher Scientific). The reactions were transformed into competent cells of *Escherichia coli* DH5α. The transformed cells were left to recover for one hour at 37°C in shaking conditions and then plated on LB plates containing the appropriate selecting antibiotic. The plasmid DNA samples were extracted and sequence-verified as described above.

### Trans-activation assays

We grew the material for protoplast-based trans-activation assays as follows. We stratified *Arabidopsis thaliana* Col-0 plants for 4 days on soil:perlite mix 1:2 (Primasta BV, Asten, The Netherlands) and then moved them to a growth chamber under short-day conditions (8 h light, 16 h dark; 20°C; 70% humidity) for a week to allow germination. The plants were then transplanted and left to grow for 3 weeks. We isolated leaf mesophyll protoplasts using the tape sandwich method (Wu *et al*., 2009). We incubated the tapes containing the tissues in an enzymatic solution [1% cellulase R-10 (Duchefa Biochemie), 0.25% macerozyme R-10 (Duchefa Biochemie), 0.4 M mannitol, 20 mM KCl, 20 mM MES pH 5.8, 10 mM CaCl2, 5 mM β-mercaptoethanol, 0.1% BSA] in the dark with gentle agitation for two hours at room temperature. We filtered the protoplasts through a 100 µm nylon mesh, washed them two times in W5 buffer (154 mM NaCl, 125 mM CaCl2, 5 mM KCl, 2 mM MES pH 5.8) and incubated them on ice for 30 minutes. We determined the protoplast number using a hemocytometer. We finally resuspended the protoplasts in MMg solution (0.4 M mannitol, 15 mM MgCl2, 5 mM MES pH 5.8). For each biological replicate, we transfected 1-5 x 10^5 protoplasts with 10 μg of pDLUC15 plasmid carrying the promoter sequence in combination with 3-5 μg of the vectors expressing the coding sequence of the transcription factors of interest by adding a PEG solution (40% PEG 4000, 0.2 M mannitol, 0.1 M CaCl2). We incubated the reactions for 5 minutes; we stopped the transfections with W5 buffer, and we washed them two times with W5 buffer. After transfection, we incubated the protoplasts for 16-18 hours at room temperature in the dark, spun them down, and removed the supernatant. We then detected the activity of the Firefly and Renilla luciferases with the Dual-luciferase reporter assay system (Promega) and the GloMax 96 Microplate Luminometer (Promega). We assayed two technical replicates for each biological replicate. The statistical analysis was done as described in the legend of the figures.

## Results

We explored the regulatory interactions that tomato exodermis-enriched TFs have with their target genes in order to understand the suberin regulatory module in exodermis. We selected three TFs, SlMYB92, SlMYB41 and SlWRKY71, as targets for this study, identified as exodermis-enriched based on tomato root gene expression profiling by Cantó-Pastor *et al*. (2024) and Kajala *et al*.(2021)(Figure 1A) and as regulators of suberin levels (SlMYB92 and SlMYB41, Cantó-Pastor *et al*., 2024). Using a translational fusion with GFP we confirmed that *SlMYB92* is expressed only in the exodermis layer of tomato hairy roots (Figure 1B, Cantó-Pastor *et al*., 2024), while exodermis-specific promoter activities of *SlMYB41* and *SlWRKY71* were previously demonstrated in Kajala *et al*. (2021).

**Figure 1:**
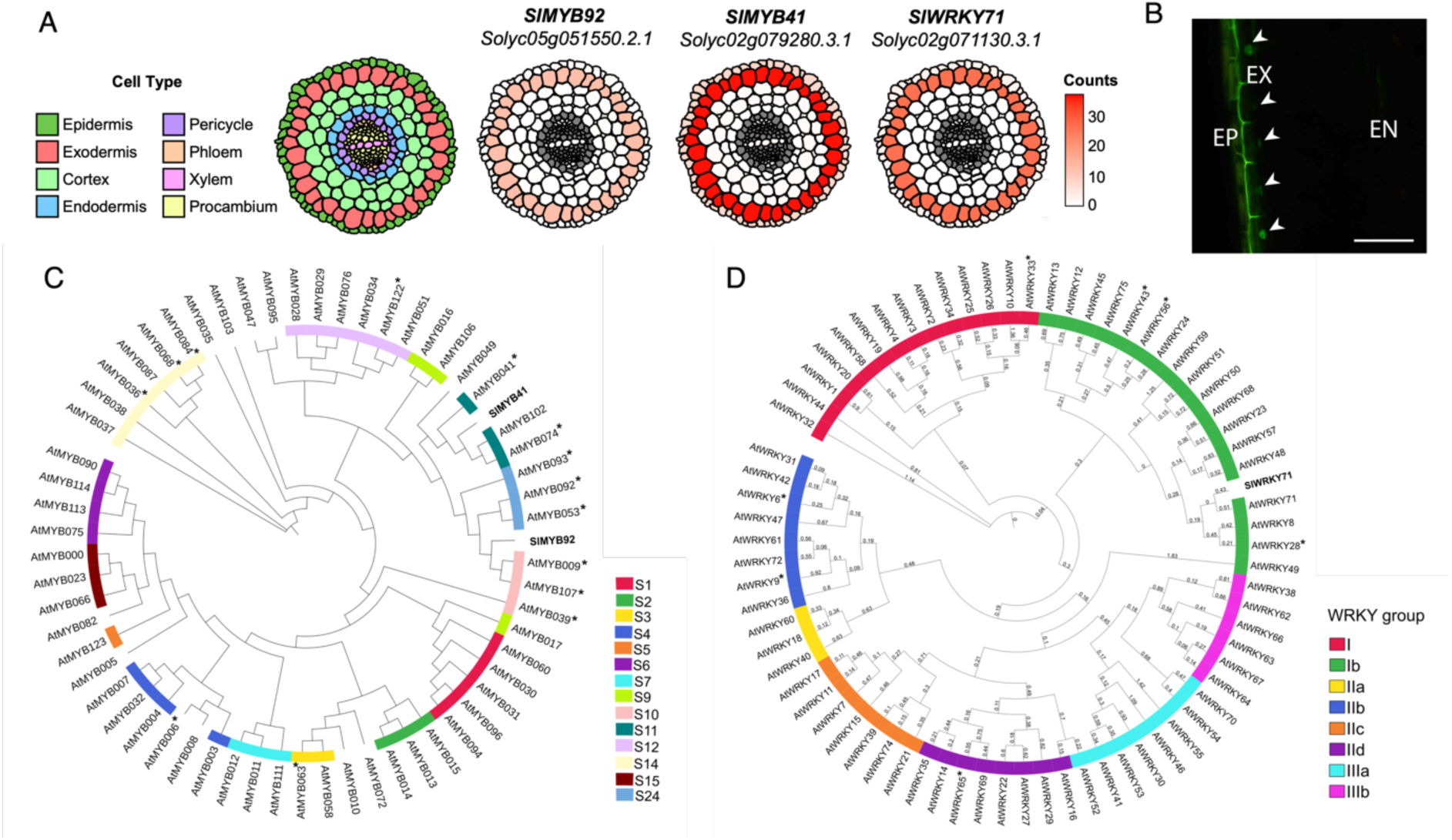
*SlMYB92*, *SlMYB41* and *SlWRKY71* are exodermis-enriched TFs with homologs with roles in regulating suberin. A: Cell type-enriched gene expression heatmaps for *SlMYB92, SlMYB41* and *SlWRKY71*of translatome data from Kajala et al. (2021). B: Transcriptional fusion *proSlMYB92:nlsGFP* drives expression in exodermis in tomato hairy root cultures. Note that autofluorescence from the exodermal and endodermal lignin are also picked up in the GFP excitation waveband. The nuclear GFP signal is indicated with arrowheads. Transcriptional fusions of *proSlMYB41 and proSlWRKY71* are previously published (Kajala *et al*., 2021). Scale bar, 100 μm. C-D: SlMYB41, SlMYB92 and SlWRKY71 are related to Arabidopsis MYB and WRKY TFs that regulate suberization. Representative phylogenetic trees showing the closest Arabidopsis homologs for (C) SlMYB4 and SlMYB92 and (D) SlWRKY71. The trees were inferred using the maximum likelihood method with 1000 bootstraps using iQ-TREE. Putative full length MYB and WRKY amino acid sequences were aligned with MAFFT. All MYBs and WRKYs indicated with an asterisk (*) have described functions linked with suberization. The MYB subgroups (“S”) are annotated in colored boxes according to Dubos et al (2010). WRKY groups are annotated with colored boxes according to Eulgem et al. (2000) and Wu et al. (2005).

The SlMYB TFs are homologous with known suberin regulators of Arabidopsis (Figure 1C). Specifically, phylogenetic analysis of *SlMYB41* and *SlMYB9*2 with the closest Arabidopsis *MYB* homologs, including 14 with described roles in suberization showed protein sequence similarity with MYB subgroup S11 (*AtMYB41, AtMYB74, AtMYB102*) and S24 (*AtMYB93*, *AtMYB92* and *AtMYB53*), respectively (Figure 1C).

For the WRKY TF, *SlWRKY71* was selected as a candidate due to its expression being induced in drought, similarly to *SlMYB92, SlMYB41*, and suberin deposition (Cantó-Pastor *et al*., 2024). Phylogenetic analysis of SlWRKY71 with homologous Arabidopsis WRKY proteins showed that SlWRKY71 is closely related to AtWRKY71, AtWRKY8 and AtWRKY28, the latter of which has previously been identified as a homolog of a candidate regulator of suberin biosynthesis in poplar (Figure 1D, Supplementary Table S1) (Rains *et al*., 2018). So, we asked if these two MYBs and SlWRKY71 form a regulatory module to coordinate the expression of suberin biosynthesis genes in the exodermis of tomato.

To identify the transcripts in the tomato root tip that are regulated by SlMYB92, SlMYB41 and SlWRKY71, we used both CRISPR/Cas9 induced knockouts and *p35S-*driven overexpression constructs for all three TFs in tomato hairy root cultures (Ron *et al*., 2014; Cantó-Pastor *et al*., 2024). Additionally, the knockout and overexpression lines show if these TFs are necessary or sufficient, respectively, for exodermal suberin deposition. For the MYB knockouts, we used our previously published *Slmyb92-*ko and *Slmyb41-*ko lines that have reduced suberization of tomato root exodermis (Cantó-Pastor *et al*., 2024). The gene editing of *SlWRKY71* generated two truncated versions of the protein *Slwrky71*-ko(4) and *Slwrky71*-ko(5) which we used in parallel in our experiments (Supplementary Figure S1). For the overexpression constructs with the near constitutive *35S* promoter, we included a C-terminal 3xFLAG tag and validated the hairy root lines by protein accumulation as detected by anti-FLAG antibody (Supplementary Figure S2). For our experiments, we selected the overexpression lines with the strongest TF accumulation (Supplementary Figure S2).

Our first question about these lines was if they had altered exodermal deposition of suberin or lignin. We set out to test the effect of the genetic perturbations by examining suberin deposition in cross-sections of the ko and OX hairy root lines and comparing them to the control (empty vector) hairy roots (Figure 2). As suberization in exodermis builds up gradually, starting with a patchy pattern and building up to a fully suberized ring, we quantified our section data accordingly. We scored whether cross-sections from individual roots had exodermis that was non-suberized (0% of cells in a section), patchy (“early-suberized”, <50% of cells in the section) or more fully suberized pattern (“late suberized”, >50% of cells in a section) (Figure 2A). For the *Slmyb92*-ko and *Slmyb41-*ko, we saw reduction in exodermal suberisation when compared to the empty vector control (Figure 2A-C), confirming that these two MYB TFs are necessary for normal suberin deposition pattern in tomato exodermis. Conversely, the overexpressors of the two *SlMYBs* had similar suberin deposition pattern as the control roots. This suggests that they are not individually sufficient to affect suberization. In *SlWRKY*-OX, we observed an absence of suberization in the exodermis (Figure 2A-C), while *Slwrky71-*kos did not affect the suberization. This indicates a repressive role for SlWRKY71 in exodermal suberin deposition with potentially other redundantly acting TFs. We then examined the lignin deposition in the exodermis of these lines, and did not observe any changes to the deposition pattern (Supplementary Figure S3), indicating that these TFs are likely to regulate only the suberin barrier deposition in exodermis.

**Figure 2:**
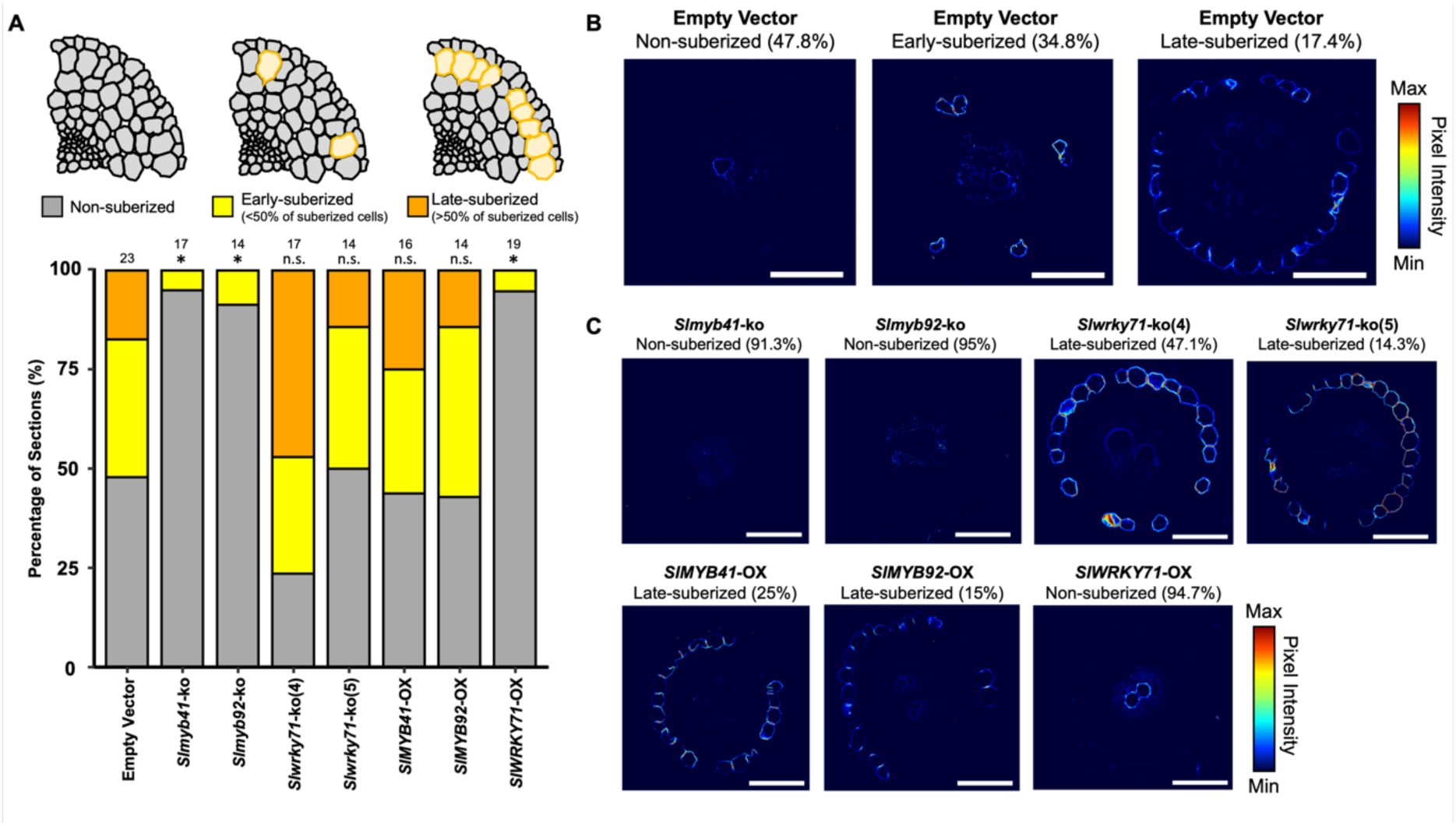
Perturbations on SlMYB41, SlMYB92 and SlWRKY71 expression affect the suberization of tomato exodermis. A: Percentage of cross sections showing a non-suberized, an early-suberized (suberin in less than 50% of exodermal cells) or a late-suberized (suberin in more than 50% of exodermal cells) exodermis phenotype in hairy-root cultures of the control (empty vector), CRISPR knock-out lines (*Slmyb41*-ko, *Slmyb92*-ko, *Slwrky71*-ko(4) and *Slwrky71*-ko(5)) and overexpressing lines (*SlMYB41*-OX, *SlMYB92*-OX and *SlWRKY71*-OX). The number above the bars represents the number (n) of individual roots examined in each genotype. Asterisks denotes statistically significant differences between each line in comparison to the empty vector control, whereas n.s. indicates no significant differences (P < 0.05, chi-squared test). B: Representative images of Fluorol Yellow-stained cross sections of the control hairy-root line (empty vector) showing a non-suberized, an early-suberized and a late-suberized phenotype. The percentage numbers on top of the images represent the percentage of cross-sections showing the indicated suberization phenotype. Scale bars represent 100 μm. C: Representative images of Fluorol Yellow-stained cross sections of the knock-out and overexpressing hairy-root lines. The percentage numbers on top of the images represent the percentage of cross-sections showing the indicated suberization phenotype. Scale bars represent 100 μm.

To identify the potential target genes of SlMYB92, SlMYB41 and SlWRKY71, we performed RNA sequencing (RNA-seq) of 1-cm root tips from hairy-root cultures of *Slmyb41*-ko, *SlMYB41*- OX, *Slmyb92*-ko, *SlMYB92*-OX, *Slwrky71*-ko(4), *Slwrky71*-ko(5), *SlWRKY71*-OX and empty vector control lines. Multidimensional scaling (MDS) analysis of the generated RNA-seq data demonstrated the reproducibility of the biological replicates for each genotype. It also revealed transcriptome similarities between the *Slmyb41*-ko and *Slmyb92*-ko lines, between the *Slwrky71*-ko(4) and *Slwrky71*-ko(5) lines, and between the *SlWRKY71*-OX and *SlMYB41*-OX lines (Figure 3A). For each line, differentially expressed genes (DEGs) were identified by comparing them to the transcriptome data of control roots with an adjusted P-value threshold of 0.05 (Figure 3B). The list of DEGs identified in all the lines are summarized in Supplementary Dataset S1. Perturbation of TF function by CRISPR-Cas9 led to notable changes in the transcriptome of hairy roots, revealing 873, 624, 828, and 869 DEGs in *Slmyb4*-ko, *Slmyb92*-ko, *Slwrky71-*ko(4), and *Slwrky71-*ko(5) knockout lines, respectively (Figure 3B). The same pattern was not observed in the overexpressing lines, where 582 genes were identified in the *SlMYB92*-OX line, whereas overexpressing *SlMYB41* and *SlWRKY71* led to the identification of a smaller number of DEGs, 165 and 126 respectively (Figure 3B). We compared the DEGs between the *Slmyb* ko lines and found that 227 (53%) out of 426 down-regulated DEGs in *Slmyb41-*ko were also down-regulated in *Slmyb92-*ko (Figure 3C). A similar pattern was observed in the list of upregulated genes in *SlMYB-*OX lines (Figure 3D), where 76 (66%) out of 115 of up-regulated genes in *SlMYB41*-OX were found to be upregulated in the *SlMYB92*-OX line (Figure 3D). The similarities observed between the DEGs in both the OX and ko lines of *SlMYB41* and *SlMYB92* indicate an overlap in the gene regulatory networks controlled by these two MYB TFs. We also compared the list of DEGs in the two independent *Slwrky71*-ko lines and found that 60% and 54% of downregulated and upregulated DEGs identified in *Slwrky71-*ko(5) line was found in the list of DEGs from *Slwrky71*- ko(4) (Figure 3D). The major overlap between the DEG lists of the two independent *Slwrky71*-ko lines indicates accurate identification of the genes regulated by this TF. For further analysis, we consider the overlapping DEGs between the independent lines as the DEGs for *Slwrky71-*ko (Figure 3D).

**Figure 3:**
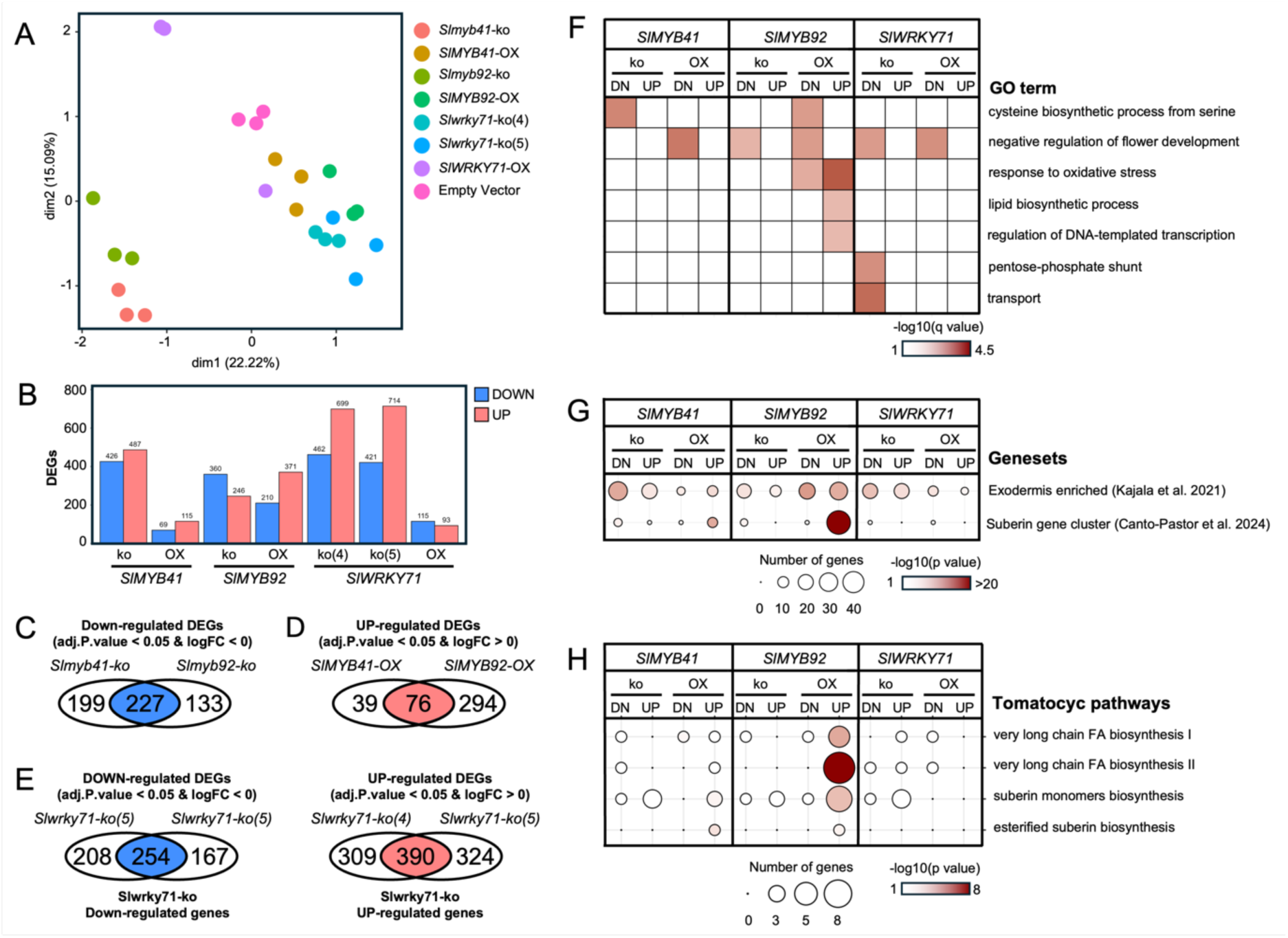
Transcriptome analysis of TF knockouts (kos) (*Slmyb41-ko, Slmyb92-ko, Slwrky71-* ko(4) *and Slwrky71-*ko(5)) and overexpressors (OX) (SlMYB41-OX, SlMYB92-OX and SlWRKY71-OX) in tomato hairy root cultures. A: Multidimensional scaling (MDS) plot showing the distance between samples. B: Number of up (FC > 0) and down (FC < 0) DEGs identified in each line with an FDR threshold of 0.05. C: Venn diagram showing the overlap between down-regulated genes in *Slmyb41-*ko and *Slmyb92-*ko. D: Venn diagrams showing the overlap between the up-regulated genes between *SlMYB41-*OX and *SlMYB92-*OX. E: Venn diagrams showing the overlap between DOWN and UP regulated DEGs between *Slwrky71*-ko(4) and *Slwrky71-*ko(5). DEGs found in both independent lines were considered as down or upregulated in *Slwrky71-*ko. F: Heatmap showing the q-value significance (-log10(q value)) of GO terms enriched in the ko and OX lists of DOWN (DN) or UP regulated list of DEGs. G: Bubble heatmap plots showing the number of DEG genes that overlap with previously annotated list of exodermis enriched (Kajala et al. 2021) and root suberin co-expressed module (Canto-Pastor et al. 2024). Statistical significance of the overlap between datasets is indicated (hypergeometric distribution). H: Bubble heatmap plots showing the number of DEG genes that overlap with genes annotated as involved with FA and suberin biosynthesis according to the tomatocyc database. Statistical significance of the overlap between datasets is indicated (hypergeometric distribution).

To identify the biological role of genes regulated by these TFs, we performed gene ontology (GO) enrichment analysis and identified the metabolism-related GO terms enriched (q value < 0.05) in the list of DEGs (Figure 3F). Lipid biosynthesis (GO:0008610) was found to be enriched in the list of upregulated (UP) genes in *SlMYB92*-OX (Figure 3F). These genes include 3- ketoacyl-CoA synthase (KCS)-encoding genes that are involved with the elongation of fatty-acids (Li-Beisson *et al*., 2010).

Interestingly, response to oxidative stress (GO:0006979) was found to be enriched in the *SlMYB92-*OX upregulated list of DEGs (Figure 3D). This is likely due to the high number of genes encoding peroxidases and catalases found in the OX lists of DEGs (Supplementary Figure S3). It has been proposed that in addition to lignin deposition, peroxidases have a role in promoting the formation of the suberin lamellae in tomato roots (Quiroga *et al*., 2000; Serra and Geldner, 2022). These results indicate that SlMYB92 is involved with the suberization of the tomato root exodermis.

To further investigate if DEGs are involved in suberin biosynthesis in the exodermis, we compared the list of DEGs to previously characterized tomato exodermis-enriched and tomato root-expressed suberin biosynthesis genes (Kajala *et al*., 2021; Cantó-Pastor *et al*., 2024) (Figure 3G). We identified a significant overlap (p < 0.01) between the DEGs in *Slmyb41-*ko*, Slmyb92-*ko*, SlMYB92-*OX*, Slwrky71-*ko(4), and *Slwrky71-*ko(5) with the previously annotated tomato exodermis-enriched genes (Kajala *et al*., 2021) (Figure 3G). Additionally, we found a significant overlap between the DEGs in *SlMYB92-*OX and *SlMYB41-*OX with the previously described list of tomato root suberin-related genes (Cantó-Pastor *et al*., 2024). Finally, to determine the extent to which these TFs regulate genes involved in the distinct steps of suberin biosynthesis, we compared the list of DEGs to the genes associated with fatty acid (FA) (very long chain FA biosynthesis I & II) and suberin biosynthesis (suberin monomers biosynthesis and esterified suberin biosynthesis) pathways in the tomato Plant Metabolic Network (PMN) (https://plantcyc.org/databases/tomatocyc/4.0). We observed a significant overlap between the DEGs in *SlMYB41-*OX and *SlMYB92-*OX with the FA and suberin-related pathways (Figure 3H). This finding suggests that MYB TFs promote the suberization of the tomato root exodermis by enhancing the expression of genes involved in the precursors and monomers of suberin biosynthesis. Notably, some of these genes encode for key enzymes of the suberin biosynthesis pathway, and enzymes such as Aliphatic Suberin Feruloyl Transferase */* ω-HYDROXYACID/FATTY ALCOHOL HYDROXY-CINNAMOYL TRANSFERASE (ASFT/FHT) (*Solyc03g097500.3.1*) (Serra and Geldner, 2022; Cantó-Pastor *et al*., 2024) and glycerol-3-phosphate acyltransferase 4 (GPAT4) (*Solyc01g094700.3.1* and other GPATs*, Solyc03g097500.3.1*, *Solyc09g014350.3.1*) (Feng *et al*., 2022; Serra and Geldner, 2022). Altogether, our results support SlMYB92 and SlMYB41 as regulators of the suberin biosynthesis program in the tomato root exodermis.

Our RNA-seq analysis also suggests an antagonistic role between the two SlMYBs and SlWRKY71 transcription factors in regulating the suberin biosynthesis program in tomato roots. For instance, 107 out of 371 upregulated genes (FDR < 0.05 and logFC > 0) in *SlMYB92-*OX were also upregulated in *Slwrky71*-ko (Figure 4A). Furthermore, similar to the peroxidase and catalase genes (Supplementary Figure S4), many suberin biosynthesis-related genes upregulated in *SlMYB92-*OX and *SlMYB41-*OX exhibited lower fold-change levels in *SlWRKY71-*OX (Figure 4B). These results suggest that these SlMYBs primarily act as positive regulators of the suberin biosynthesis program in the tomato root exodermis, while SlWRKY71 functions as a repressor of exodermis suberization. Moreover, these results indicate that these TFs may balance each other’s actions to fine-tune the expression of suberin biosynthesis genes in the tomato root exodermis.

**Figure 4:**
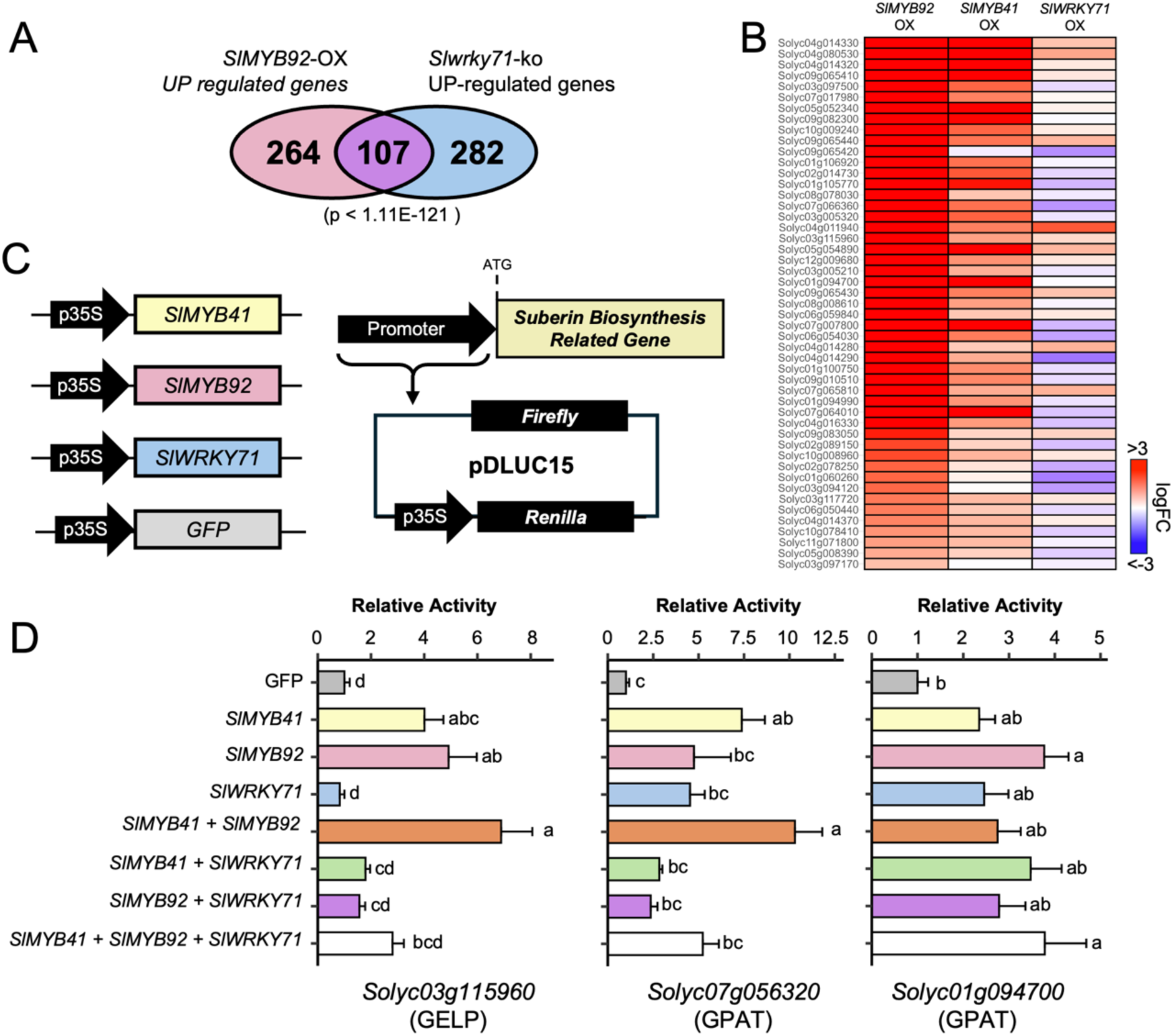
SlMYBs and SlWRKY71 antagonistically regulate the expression of suberin related biosynthesis genes. A: Venn diagram showing the overlap between *SlMYB92-*OX, and *Slwrky71-*ko upregulated genes (FDR < 0.05, logFC > 0). Statistical significance of the overlap between genesets is indicated (hypergeometric distribution). B: Heatmap showing the logFC values of genes in the *SlMYB92-*OX*, SlMYB41-*OX and *SlWRKY71-* OX. Values are presented for the genes that are significantly up-regulated in the *SlMYB92-OX* and annotated in the suberin co-expression cluster (Canto-Pastor et al. 2024). C: Schematic diagrams of constructs used for the transactivation assays. D: Transactivation assays in protoplasts of *A. thaliana*. The protoplasts were co-transformed with the pDLUC15 vector expressing the Firefly luciferase under the control of the promoter of interest and with different combinations of overexpressed transcription factors, as indicated in the X-axis. The pDLUC15 contained also the *35S::Renilla* luciferase, used as internal control. The Firefly to Renilla luciferase activity ratios were normalized to the average ratios of the GFP samples. Data represent mean ± SE (n = 4-5). Different letters indicate statistically significant differences (1-way ANOVA followed by a post-hoc Tukey test, p < 0.05).

To further test the function of these TFs in the transcription of their target genes, we performed transactivation assays in Arabidopsis mesophyll protoplasts using individual TFs and all their combinations. A construct overexpressing GFP was used to establish a baseline for the assay. A schematization of the constructs used is shown in Figure 4C. For these assays we tested the promoter regions of the two *SlMYBs* and of *SlWRKY71,* to see whether they regulate their own expression, in addition to three other candidate genes selected for their involvement in the suberin biosynthesis pathway and their transcriptional response in the transgenic hairy roots (Supplementary Figure S4). Among the different promoter regions used, only *Solyc03g115960.3.1* (*GDSL esterase/lipase38* (*SlGELP38*)), *Solyc07g056320.4.1* (*Glycerol-3- phosphate acyltransferase*) and *Solyc01g094700.5.1* (*SlGPAT4*) showed a positive transactivation (Figure 4D, Supplementary Figure S5). Specifically, *SlGPAT4* was transactivated by SlMYB92 alone and when in combination with the other two TFs. On the other hand, *SlGELP38* and *Solyc07g056320.4.1* were transactivated by both SlMYB92 and SlMYB41 and their combination, but not by SlWRKY71 (Figure 4D). When SlWRKY71 was combined with either one or with both MYBs, it seemed to act like a repressor of promoter activity (Figure 4D). This suggests that SlMYB92 and SlMYB41 are positive regulators of these genes while SlWRKY71 counteracts the function of the two SlMYBs (Figure 4). Overall, it appears that the MYBs are direct activators of promoters of suberin-related genes and their effect may be additive, while SlWRKY71 can act as an activator or an inhibitor of promoter activity depending on the promoter and context of other TFs present.

## Discussion

### SlMYB92 and SlMYB41 are positive regulators of the suberin biosynthesis program in tomato root exodermis

In Arabidopsis roots, suberin accumulates in the endodermal cell layer, whereas in tomato, suberization occurs in the exodermal cell layer. Despite this spatial distinction, it has been shown that both processes are controlled by similar components (Cantó-Pastor *et al*., 2024). Several reports have shown the role of MYB TFs in the control of the suberization in the root endodermal cell layer. In Arabidopsis, several MYB TFs, including AtMYB41, AtMYB53, AtMYB92, and AtMYB93, have been shown to be involved in the biosynthesis of suberin in the root endodermal cell layer (Shukla *et al*., 2021). Additionally, 13 key MYB TFs were identified in transcriptional regulatory network to control the suberization and lignification in the Arabidopsis endodermis (Xu *et al*., 2022b). In this sense, it is likely that MYB TFs would also play an important role in the suberization of the root exodermis in tomato. Here we provide several lines of evidence that show that SlMYB92 and SlMYB41 can act as positive regulators of suberin biosynthesis by regulating the expression of several genes involved with the different steps of suberin biosynthesis in tomato roots. First, as previously reported, *SlMYB92* and *SlMYB41* are expressed primarily at the exodermal cell layer of the tomato root (Figure 1) (Kajala *et al*., 2021; Cantó-Pastor *et al*., 2024). Second, perturbations of *SlMYB92* and *SlMYB41* coding sequences by CRISPR/Cas9 resulted in a significant delay in suberin deposition in the tomato root exodermis (Figure 2, Cantó-Pastor *et al*., 2024). Third, overexpression of *SlMYB92* and *SlMYB41* in tomato hairy root cultures resulted in the upregulation of several genes involved with suberin biosynthesis (Supplementary Dataset S2). However, we did not observe increased accumulation of suberin in the *SlMYB*-OX lines, indicating that they might work together to upregulate the complete suberin biosynthesis and deposition pathways. Fourth, we showed that promoters of suberin biosynthesis genes are transactivated by SlMYB92 and SlMYB41 in Arabidopsis leaf protoplasts (Figure 4D), indicating direct activation of the promoter activity. Together, our results provide strong evidence that SlMYB92 and SlMYB41 act as positive regulators of suberin biosynthesis in the tomato root exodermis.

Given the challenges of generating transgenic tomato plants, our genetic perturbations were done with the tomato hairy root cultures and transactivation assays in Arabidopsis leaf protoplasts, which introduces the caveats of using model systems that are different from the tomato embryonic roots. We observed lower levels and higher variation of exodermal suberization in the tomato hairy root cultures compared to primary roots (Kajala *et al*., 2021; Cantó-Pastor *et al*., 2024). We did not observe *SlMYB* overexpressors driving increased suberization unlike previously observed for other species (Kosma *et al*., 2014; Shukla *et al*., 2021; Chen *et al*., 2024). This could be due to inadvertent selection towards weakly suberizing alleles. If the SlMYBs were sufficient to drive suberization, their effect expressed under the near-constitutive 35S promoter could lead to inhibition of root growth, and hence not being selected for hairy root cultures. Additionally, the heterologous trans-activation assay might be affected by the factors present in the heterologous tissue of a different species. While these challenges may affect the interpretation of the results, the different lines of inquiry support each other and are consistent with the Arabidopsis literature for the endodermis.

### SlMYB92 and SlMYB41 regulate genes involved in suberin biosynthesis and deposition

Suberin is a highly heterogeneous biopolymer comprised of a variety of aliphatic long chain fatty acids and their oxidized derivatives, glycerol and ferulic acid (Shukla and Barberon, 2021; Serra and Geldner, 2022). Due to its biochemical complexity, the action of several enzymes that operate in distinct compartments of the cell needs to be highly coordinated. Here, we show that SlMYB92 and SlMYB41 can regulate the expression of several genes involved in the many steps required for the biosynthesis of suberin monomers, their transport and polymerization in the apoplast (summarized in Supplementary Dataset S2). The precursors of suberin monomers are produced via two major biochemical pathways: very-long-chain fatty acid (VLCFA) and phenylpropanoid biosynthesis pathways (Shukla and Barberon, 2021). The fatty-acid elongase (FAE) complex catalyzes the elongation of acyl-CoA into VLCFA. Within this complex, the ketoacyl-CoA synthases (KCS) enzyme carries out the rate-limiting step (Haslam and Kunst, 2013; Batsale *et al*., 2021; Serra and Geldner, 2022). Here we showed that *SlKCS1* (*Solyc10g009240.3.1*), *SlKCS6* (*Solyc02g085870.3.1*) and *SlKCS20* (*Solyc03g005320.3.1* and *Solyc09g083050.3.1*) were up-regulated in the *SlMYB92*-OX line while *SlKCS11* (*Solyc08g067410.2.1*) was down-regulated in *Slmyb41-*ko and *Slwrky71-*ko lines. These VLFAs are further modified by laccases (LACSs) and CYTOCHROME P450s (CYPs) (Serra and Geldner, 2022). We found three LACS encoding genes to be upregulated in *SlMYB92*-OX (*Solyc04g011900.4.1, Solyc05g052340.4.1, Solyc06g082240.2.1/SlLAC3-like*) and one downregulated in *Slmyb41-*ko (*Solyc06g050530.3.1/SlLAC12-like*). Notably, tomato orthologues of *CYP86A1* (*Solyc06g076800.3.1*) and *CYP86B1* (*Solyc02g014730.3.1*), two main CYPs involved with the ω-oxidation of FAs and endodermis suberization in Arabidopsis (Li *et al*., 2007; Höfer *et al*., 2008; Compagnon *et al*., 2009) were upregulated in hairy-roots overexpressing SlMYB92 (Supplementary Dataset S2). Knock-out mutations in the latter (SlCYP86B1) resulted in a reduction of exodermis suberization in tomato hairy roots (Cantó-Pastor *et al*., 2024).

In the final steps of suberin monomer biosynthesis, a feruloyl transferase (ASFT/FHT) and GPATs are involved with the feruloylation and glyceration of suberin precursors, respectively (Serra and Geldner, 2022). It has been previously shown that knocking down *SlASFT* and *SlGPAT4* in tomato hairy roots resulted in dramatic reduction on exodermis suberization (Cantó-Pastor *et al*., 2024). We found that *SlASFT* (*Solyc03g097500.3.1*) and *SlGPAT6* (*Solyc03g097500.3.1*) to be upregulated in *SlMYB92*-OX while *SlGPAT4* (*Solyc01g094700.5.1*) was found upregulated in *SlMYB92*-OX and *SlMYB41*-OX (Supplementary Dataset S2). The transport of suberin monomers to the apoplast is mediated by lipid transfer proteins (LTPs) and ATP-binding cassette transporters of the subfamily G (ABCG) (Shukla and Barberon, 2021). In Arabidopsis, AtLTPg15 was found to be important for the transport of suberin monomers export to the seed coat (Lee and Suh, 2018). Interestingly, we observed that the tomato orthologue of *AtLTPg15* (*Solyc09g065430.4.1*) was upregulated by overexpressing *SlMYB92* and *SlMYB41* (Supplementary Dataset S2). Additionally, two ABCG-encoding genes (*Solyc03g019760.4.1* and *Solyc05g054890.4.1/SlABCG2*) were upregulated in *SlMYB92*-OX, and another *ABCG* gene (*Solyc05g051530.5.1*) was downregulated in *Slmyb92-*ko and *Slmyb41-*ko but upregulated in *SlMYB92*-OX compared to the control hairy root culture.

After the transport of suberin monomers, the GDSL-type Esterase/Lipase Protein (GELP) family are involved with the suberin polymerization in the apoplastic space. A recent study has shown that quintuple *GELP* mutants (*gelp22-38-49-51-96*) have drastic reduction in endodermis suberization (Ursache *et al*., 2021). We observed that a *GELP* gene (*Solyc03g115960.3.1/SlGELP38*) previously identified as specifically expressed in the exodermal cell layer (Kajala *et al*., 2021; Cantó-Pastor *et al*., 2024) was upregulated by the overexpression of *SlMYB92* (Supplementary Dataset S2). Additionally, four other GELP-encoding genes (*Solyc02g070610.3.1, Solyc10g085170.3.1, Solyc11g051060.2.1, Solyc10g076740.3.1*) were downregulated in the *Slmyb41*-ko or *Slmyb92*-ko (Supplementary Dataset S2).

Altogether, these results show that SlMYB92 and SlMYB41 are major regulators of exodermis suberization by promoting the expression of genes involved with the biosynthesis, transport and polymerization of suberin in tomato roots.

### The role of MYBs in gene regulation

MYBs are among the largest families of TFs discovered in plants, extensively characterized for their role in regulating responses to various abiotic stresses, such as drought and high salinity. Additionally, they play a pivotal role in regulating the biosynthetic pathways of secondary metabolites, including anthocyanins, proanthocyanidins, flavonols, lignin, and suberin (Cao *et al*., 2020; Wang *et al*., 2021; Xu *et al*., 2022b). MYBs can carry out their regulatory functions as monomers, homodimers, heterodimers or as part of protein complexes (Pireyre and Burow, 2015). For example, in the anthocyanin and proanthocyanidin pathways, MYBs interact with bHLH TFs, and, together with a WD40 protein, form the MBW complex, which regulates the transcription of biosynthetic genes (Xu *et al*., 2015). MYBs that participate in the MBW complex contain a bHLH-binding motif in their sequence, [DE]Lx2[RK]x3Lx6Lx3R, responsible for the interaction between MYBs and bHLH (Zimmermann *et al*., 2004). The MYB sequences studied here do not contain this motif, suggesting that SlMYB92 and SlMYB41 do not require an MBW complex to activate transcription. In fact, the suberin biosynthetic pathway is primarily regulated by MYB, NAC, and WRKY TFs (Woolfson *et al*., 2022). This observation aligns with our evidence that the two MYBs examined in this study can transactivate gene expression independently of bHLH transcription factors (Figure 4D).

The primary role of both MYBs seems to be regulating the expression of biosynthetic genes, with minimal impact on the regulation of other TFs (Supplementary Dataset S2). For instance, overexpression of *SlMYB41* leads to the upregulation of only one WRKY TF (*Solyc03g095770.3.1*, *AtWRKY70*), while *SlMYB92* overexpression results in the upregulation of just two WRKY TFs (*Solyc01g095630.3.1* and *Solyc09g015770.3.1)*. Notably, *SlMYB41* overexpression does not affect the upregulation of any MYB-encoding gene, whereas *SlMYB92* overexpression induces the upregulation of four MYBs (*Solyc03g093930.5.1*, *Solyc06g075660.4.1*, *Solyc06g065100.3.1*, *Solyc02g082040.3.1*). Additionally, in the *Slmyb41* and *Slmyb92* ko lines, only three MYBs were found to be downregulated (*Solyc05g007160.3.1*, *Solyc02g088190.5.1*, *Solyc09g090790.3.1*). This is similar to the multi-hierarchical regulatory network for Arabidopsis endodermal suberin and lignin proposed by Xu *et al*., (2022b), where AtMYB92 and AtMYB41 are in separate tiers and distinct branches of the network. Conversely to our data, in the network by Xu and colleagues, AtMYB41 is more of a hub coordinator and would be expected to induce higher number of DEGs when mutated or induced, while AtMYB92 is on the last tier, with low impact on other TFs. It is likely that the way in which suberin GRN is connected differs between species, cells and tissues, even if they utilize orthologous MYB TFs.

Moreover, the TFs studied here do not exhibit self-activation and do not appear to regulate each other’s expression (Supplementary Figure S4). This suggests that other MYBs may be responsible for their regulation. Additionally, given that the ko lines of both *SlMYB41* and *SlMYB92* had reduced suberin (Cantó-Pastor *et al*., 2024), it indicates that the *SlMYBs* are working as necessary but not directly connected nodes in an exodermal suberin GRN. Alternatively, it is possible that some of these regulatory interactions are dependent on e.g. having the right ratio of binding partners in a MYB heterodimer. Further studies are required to elucidate the regulatory mechanisms upstream the TFs investigated in this study.

### Antagonistic interaction of MYBs and WRKYs in regulating suberin

Together with MYBs, WRKYs appear as one of the most relevant TF families involved in suberin biosynthesis in several species and tissue types (Lashbrooke *et al*., 2016). WRKYs modulate gene expression by binding to W-box *cis*-regulatory elements of stress-induced genes, and control mainly senescence, stress and defense responses (Eulgem *et al*., 2000; Rushton *et al*., 2010).

Expression of a WRKY TF in rice roots (LOC_Os01g53260), together with MYBs and other TFs, have been shown to enhanced radial oxygen loss (ROL) barrier formation (including suberin and lignin), and WRKY *cis*-regulatory elements were enriched in the promoters of the majority of upregulated genes in this tissue (Shiono *et al*., 2014). Arabidopsis MYB107 was found to be a positive regulator of suberin biosynthesis genes during seed development, and WRKYs were among the TFs that were either co-expressed with *MYB107* or co-supressed in *myb107* (Gou *et al*., 2017). Among the co-expressed partners were *AtWRKY56* (*At1g64000*) and *AtWRKY43* (*At2g46130*) (Supplementary Table S1, Gou *et al*., 2017). Furthermore, WRKYs were listed as candidate for suberin biosynthesis and cork regulation in cork oak (*Quercus suber*) together with MYBs, and a homolog of *AtWRKY43* was among the candidate with high fold change values (Soler *et al*., 2007). Homologs of *AtWRKY65*, *AtWRKY43*, *AtWRKY6* and *AtWRKY28* were also among the transcripts with the highest fold change in poplar bark, being potential candidates for biosynthesis and regulation of poplar suberin (Rains *et al*., 2018).

The WRKY and MYB TFs interact also in defense response mechanisms in a complex way, exhibiting both positive and negative interactions. Barco and Clay (2019) observed two overlapping regulatory functions for AtWRKY33 and AtMYB51 in the expression of defense-responsive specialize metabolism genes: while AtWRKY33 activated and AtMYB51 repressed the expression of one biosynthetic enzyme (CYP82C2), both TFs activate the expression of other two enzymes (CYP79B2 and CYP79B3). According to the authors, this indicates a hierarchical TF cascade were AtWRKY33 function as the condition-dependent master regulator and AtMYB51 as the dual functional regulator (Barco and Clay, 2019). Interestingly, the authors also found that AtWRKY33 and AtMYB51 do not co-localize on the promoters of these enzymes, suggesting that these TFs likely alternate in binding the promoter regions.

There are also examples of WRKY and MYB TFs working together as an activating complex. One such example is the transcription factor VqWRKY53 in Chinese wild grape (*Vitis quinquangularis*), induced by pathogen infection, forms a transcriptional complex with VqMYB14 and VqMYB15. This complex positively regulates stilbene accumulation, a phytoalexin produced in response to abiotic and biotic stresses, thereby contributing to disease resistance (Wang *et al*., 2020a).

Here, we have uncovered an antagonistic interaction between inhibitory SlWRKY71 and activating SlMYB41 and SlMYB92. The genetic perturbations of *SlWRKY71* indicated that SlWRKY71 is a repressor of tomato exodermal suberin (Figure 2), while SlMYB92 and SlMYB41 are necessary for tomato exodermal suberin deposition (Cantó-Pastor *et al*., 2024). The lack of statistically significant phenotype for *SlWRKY71-*kos may be explained either the mutation having a subtle effect on the suberin pattern at the sampled location, or by other, redundant TFs having the same antagonistic role.

Furthermore, we showed that these TFs antagonistically regulate expression of suberin-related genes in tomato exodermis (Figure 4). We observed an overlap of DEGs induced by *SlMYB41* and *SlMYB92* overexpression and *Slwrky71* knockout, indicating that the same target genes are activated by SlMYBs as are repressed by SlWRKY71 (Figure 4A). We also observed that the *SlWRKY71*-OX DEGs had a contrasting behavior to the *SlMYB*-OX lines (Figure 4B). Finally, we showed that the presence of SlWRKY71 with the two SlMYBs prevents their ability to induce promoter activity in Arabidopsis protoplasts for two target promoters (Figure 4D).

The potential mechanisms of this MYB-WRKY antagonism could be by interactions at different levels. As described in the examples above, it could arise from direct MYB-WRKY binding, or by competitive binding of WRKY and the MYB TFs on the target promoters. Additionally, SlWRKY71 mechanism of repression can also be amplified through the repression of other WRKY TFs. For instance, we found that *SlWRKY41* and *SlWRKY70* were up-regulated in *Slwrky71*-ko lines (Supplementary Dataset S2). These two SlWRKY TFs were found to be upregulated in the SlMYB overexpressing lines (Supplementary Dataset S2). This suggests that there may be downstream TFs regulated antagonistically by SlWRKY71 and the two SlMYBs.

Interestingly, *SlWRKY71* expression is drought-induced (Cantó-Pastor *et al*., 2024) alongside with the *SlMYBs* and suberin deposition. Why would both activators and repressors of the suberin gene expression be induced simultaneously in tomato exodermis? This might represent a pre-set pulse in suberin biosynthetic gene expression to ensure that available resources are not overspent on making too much suberin. These nuances arising from antagonistic interactions in the exodermis suberin GRNs may set a complex challenge for endeavours of manipulating suberin levels for improved drought resilience and sequestration of carbon dioxide from the atmosphere into suberin. The future challenges involve understanding the importance of the repressors of suberin in plant fitness and characterizing the network more fully.

## Abbreviations

ABCG: ATP-binding cassette transporters of the subfamily G
ASFT: Aliphatic suberin feruloyl transferase
BF: Basic fuchsin
CW: Calcofluor white
CYP: CYTOCHROME P450
DEG: Differentially Expressed Gene
FA: Fatty Acid
FY: Fluorol yellow
GELP: GDSL-type Esterase/Lipase Protein
GO: Gene ontology
GPAT: Glycerol-3-phosphate acyltransferase
GRN: Gene regulatory network
KCS: Ketoacyl-CoA synthase
LACS: Laccase
LTP: Lipid transfer protein
ko: knock out
MDS: Multidimensional scaling
MYB: Myeoloblastosis
OX: Overexpressor
TF: Transcription factor
VLCFA: Very-long-chain fatty acid

## Supplementary files

Dataset S1: Differentially Expressed Genes (DEGs) in tomato hairy-root cultures.

Dataset S2: Combined summary of DEGs.

Table S1: *MYB41*, *MYB92* and *WRKY71* homologs from Arabidopsis and summary of their potential roles in regulating suberization.

Table S2: Primers used to create the overexpression constructs.

Table S3: guide RNAs (gRNAs) designed to knock out *SlWRKY71*.

Table S4: List of promoters used in trans-activation assays.

Table S5: Primers used to amplify the promoters used for transactivation assays.

Table S6: Genomic sequences of promoters cloned for transactivation assays.

Table S7: Primers used to amplify coding regions of *SlMYB92*, *SlMYB41* and *SlWRKY71* for transactivation assays.

## Acknowledgements

We thank Tuğba Akyüz, Chrysa Pantazopoulou, Linge Li, Gijs van Asselt, Chenyun Hsieh, and Alise Žvigule for experimental assistance, and Kevin Morimoto for critical reading of the manuscript. The M82 seeds were provided by C.M. Rick Tomato Genetics Resource Center.

## Author contributions

KK, SMB: Conceptualization, Supervision, and Funding Acquisition LJ, SB, MASA, RK, ACP: Investigation LJ, SB, MASA, RK: Data Curation, Formal Analysis, Visualization. KK, LJ, SB, MASA, RK: Writing – Original Draft.

All authors: Writing – Review & Editing.

All authors read and have approved the final manuscript.

## Data availability

The RNA sequencing data from this study are openly available in NCBI GEO repository reference number GSE278561.

---

**Figure S1:**
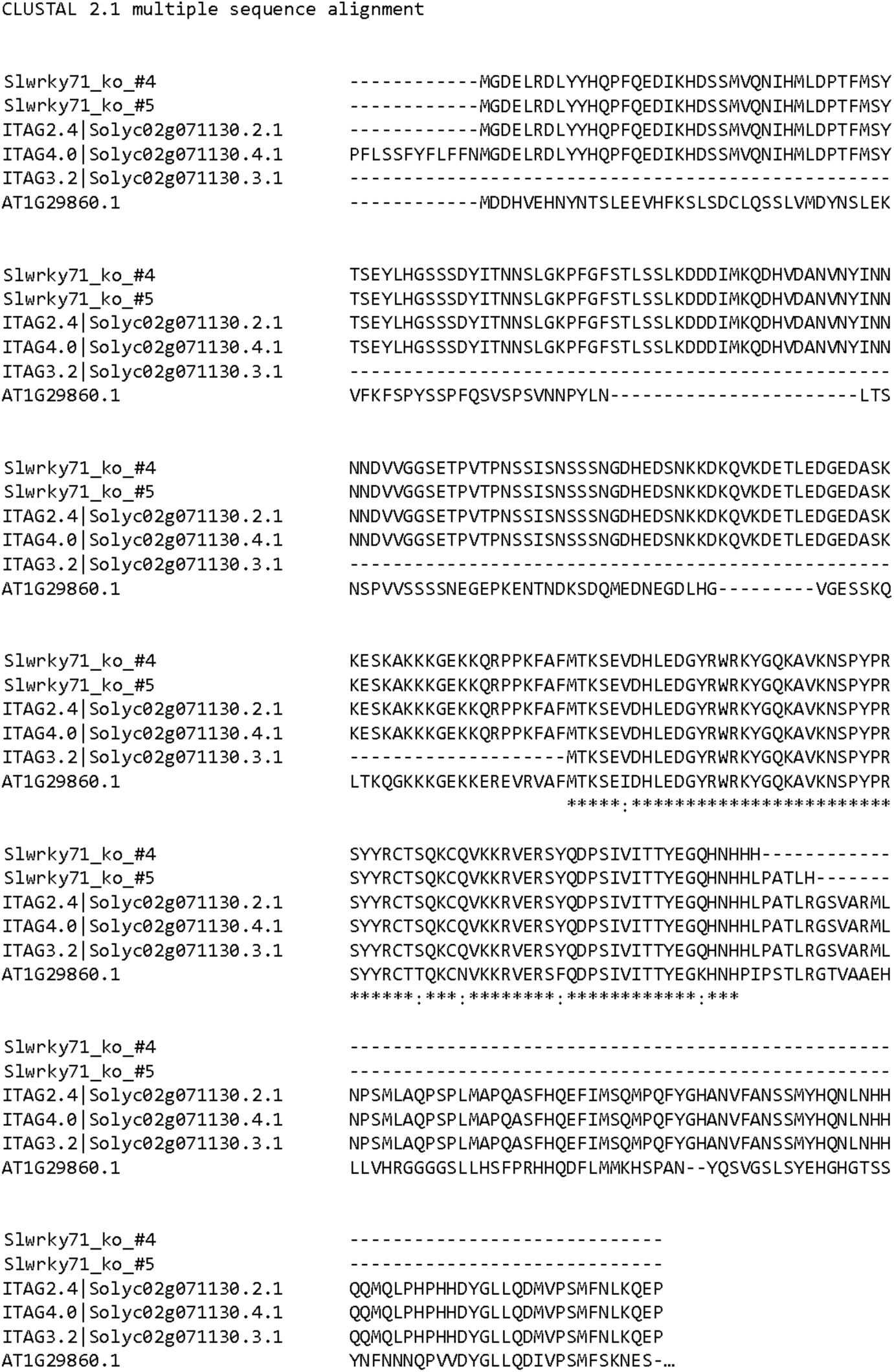
SlWRKY71 sequences and truncations generated with gene editing. Multiple sequence alignment made using ClustalW. Three different versions of SlWRKY71 from three tomato genome annotations (ITAG2.4, ITAG3.2 and ITAG4.0) were obtained from Phytozome, while the one corresponding to *A. thaliana* AT1G29860.1 was obtained from TAIR Araport 11. The protein sequences of *Slwrky71*-ko(4) and *Slwrky71*-ko(5) lines were extrapolated based on the sequencing results of the genomic fragment.

**Figure S2:**
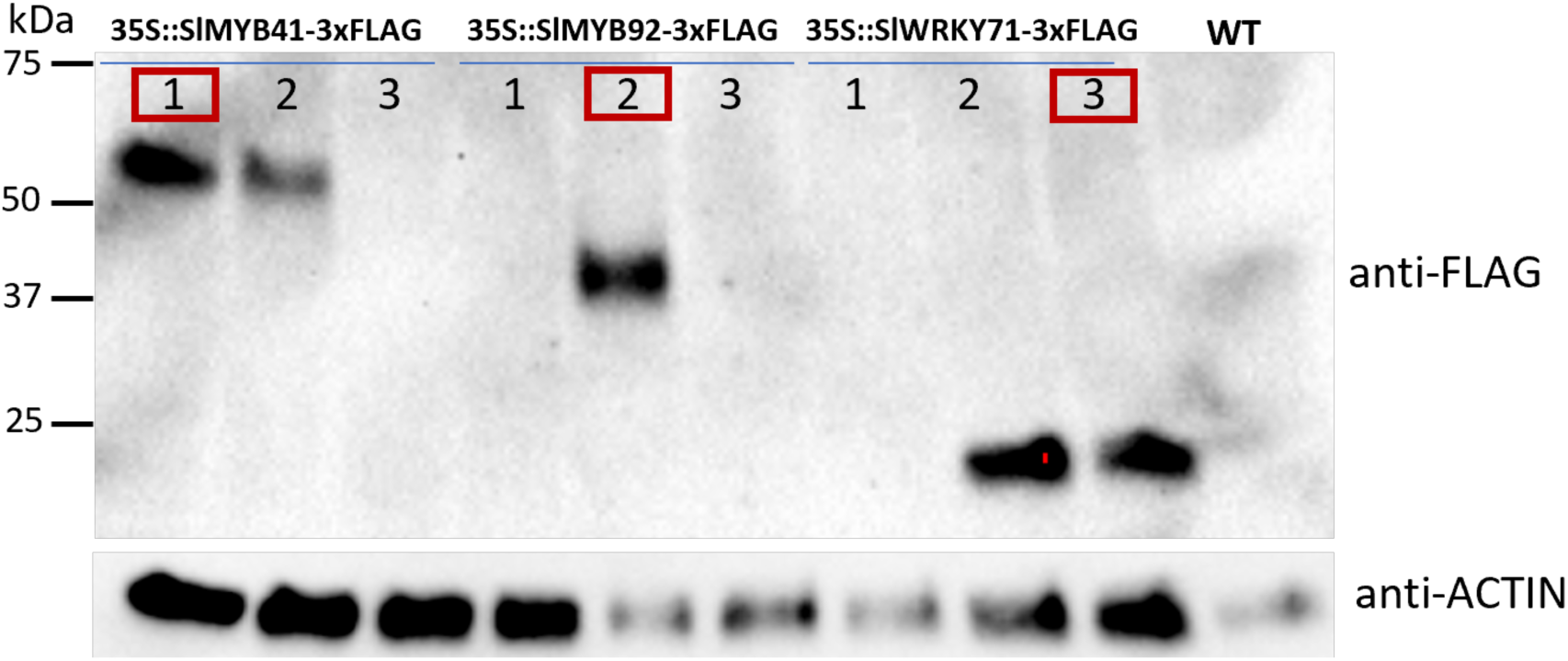
TF overexpression is detected on protein level. Western blot experiment using total protein lysates of hairy roots. Three independent transgenic lines were used to verify the presence of the overexpressed proteins with the use of the anti-FLAG antibody. The anti-ACTIN antibody was instead used to verify the correct loading of the samples. One line for each construct (circled in red) was selected for further experiments.

**Figure S3:**
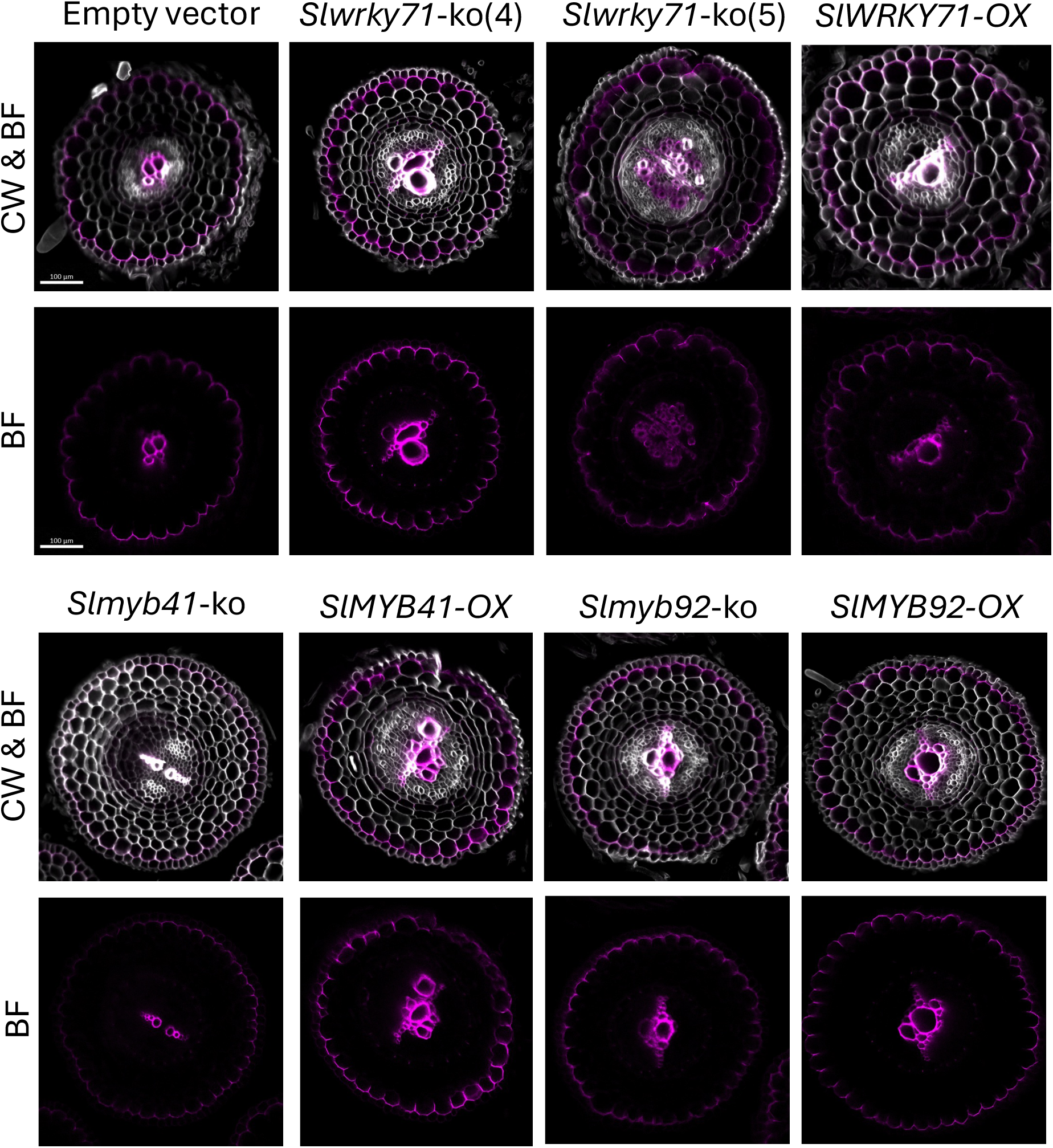
Exodermal lignin is unaffected by *SlMYB92*, *SlMYB41* and *SlWRKY71* perturbations. Representative images of root cross sections stained with Basic Fuchsin (BF, represented in magenta) and Calcofluor White (CW, represented in white). Lignin in the exodermal cell file remains unaffected for each genotype. Scale bars represent 100 μm.

**Figure S4:**
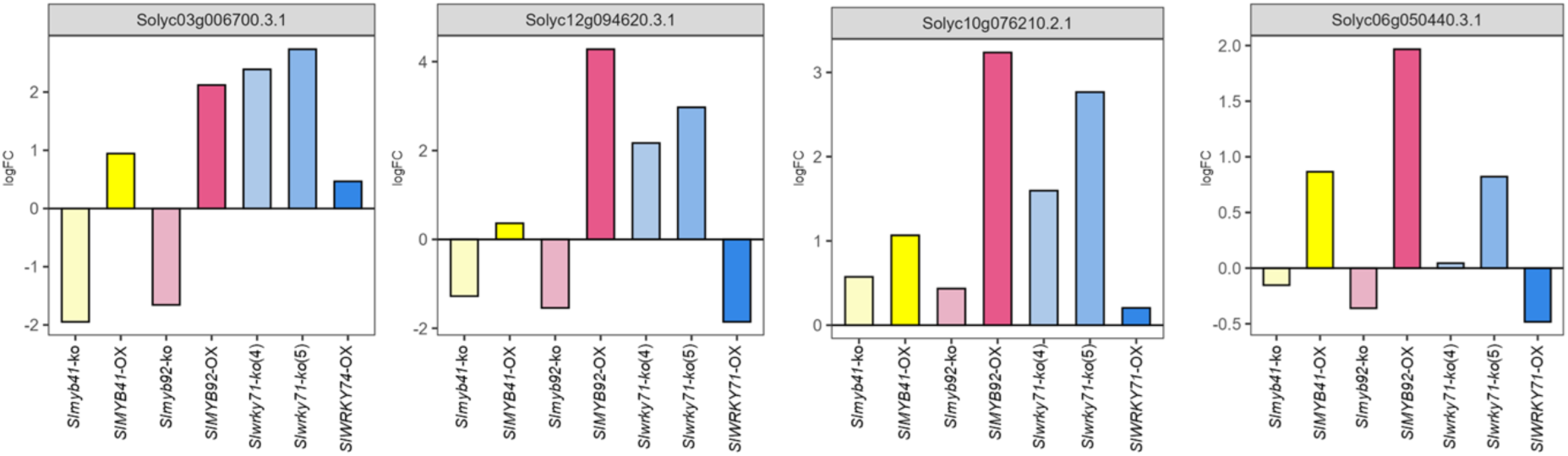
Expression of peroxidases and catalases is affected in TF knock outs and overexpressors. logFC of peroxidases and catalases in *Slmyb41-*ko, *SlMYB41-*OX, *Slmyb92-*ko, *SlMYB92-*OX, *Slwrky71-*ko(4), *Slwrky71-*ko(5) and SlWRKY71-OX in comparison to WT hairy-roots.

**Figure S5:**
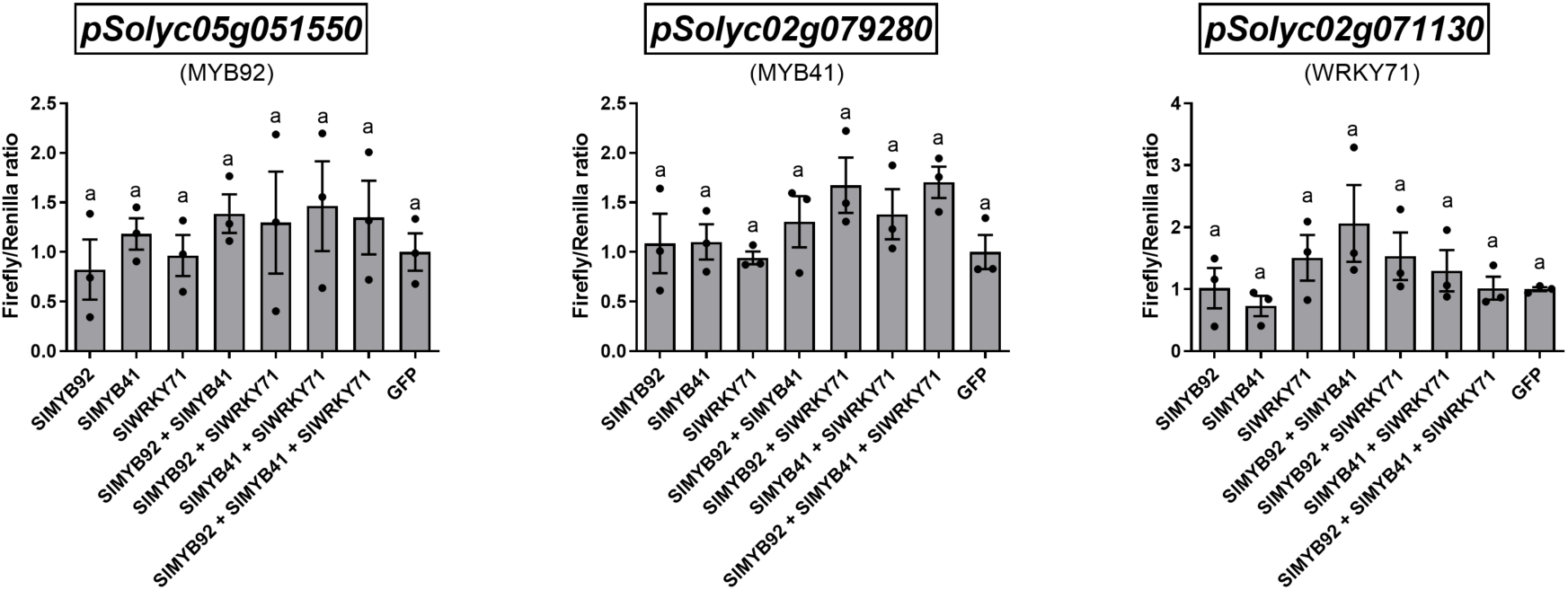
Transactivation assays of *SlMYB92*, *SlMYB41* and *SlWRKY71* promoters in protoplasts of *A. thaliana*. The protoplasts were co-transformed with the pDLUC15 vector expressing the firefly luciferase under the control of the promoter of interest and with different combinations of overexpressed transcription factors, as indicated in the X axis. The pDLUC15 contained also the *35S::Renilla* luciferase, used as internal control. Instead, when the transactivation of *pSolyc02g071130* in the vector pGWL7 was investigated, the samples were also transfected with a separate vector containing the *35S::Renilla* luciferase. The Firefly to Renilla luciferase activity ratios were normalized to the average ratios of the GFP samples. Data represents mean ± SE, each dot represents a biological replicate (n = 3). Different letters indicate statistically significant differences (1-way ANOVA followed by a post-hoc Tukey test, p < 0.05).

